## Supplemental File for "Mutation and recombination parameters in zebra finch are similar to those in mammals"

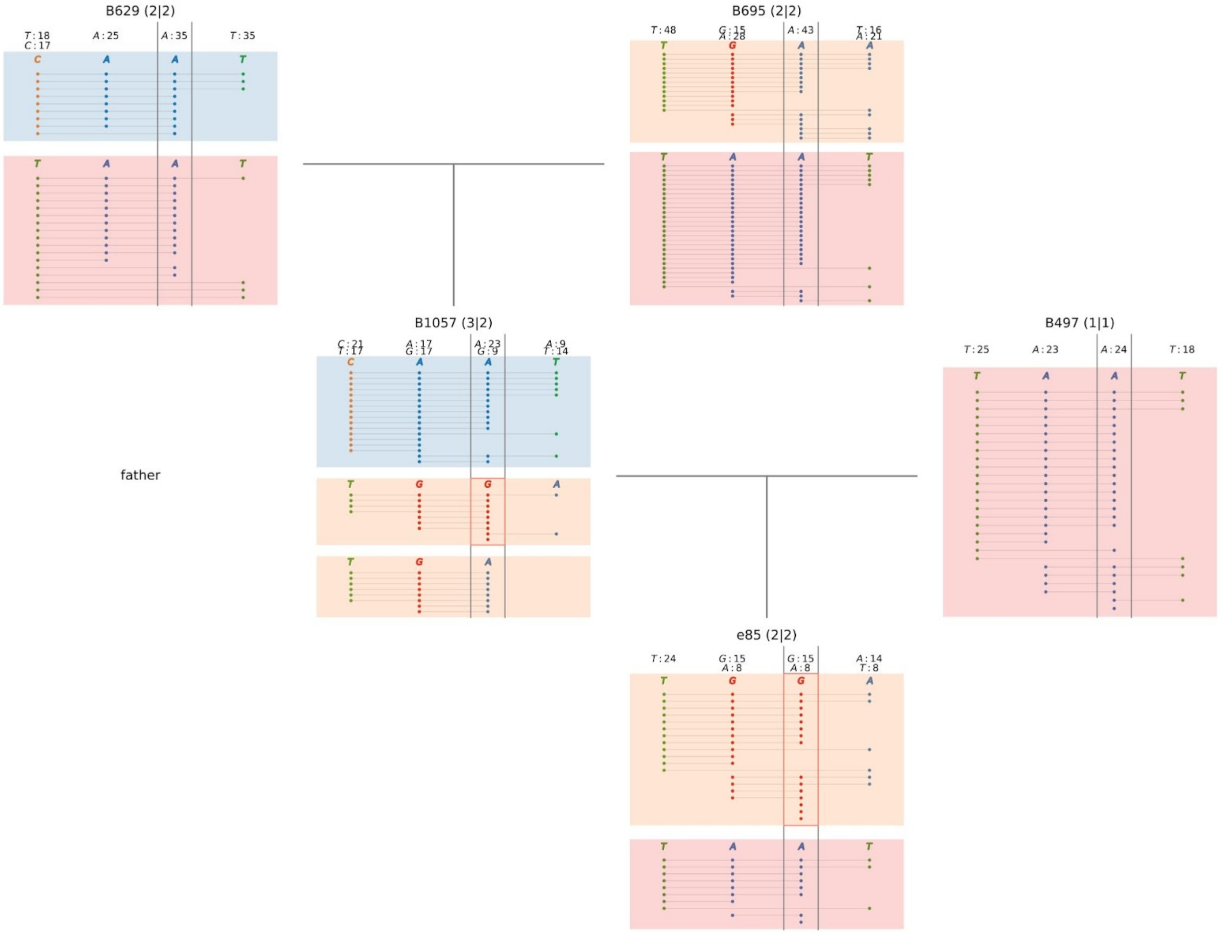


**Figure S1. Example of a DNM that likely occurred during the development of the proband (B1057, central panel).** For each of the five individuals in this pedigree, the nucleotides (circles) carried by pairs of sequencing reads spanning multiple informative heterozygous sites (horizontal lines) are shown. The mutated position is indicated by gray vertical lines, and colors represent groups of reads supporting the same haplotype (excluding the mutated position). In haplotypes carrying the DNM, the mutated position is highlighted by a red rectangle. In carrier individuals, all sequencing reads are represented for the mutated position (regardless of whether they span multiple informative sites). In the first generation (parents of the proband), the mother is positioned on the left. The annotation left to the panel of the proband shows the parental haplotype carrying the DNM. Each panel indicates the number of reads supporting each allele (top side), regardless of whether they span more than one informative site.

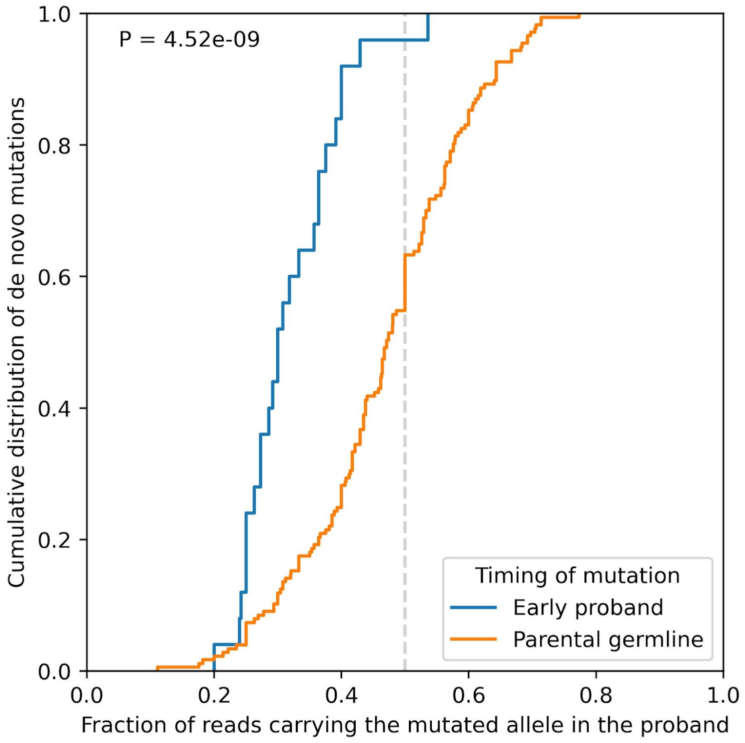


**Figure S2. Cumulative distribution of the fraction of sequencing reads supporting the mutated allele.** Cumulative distribution of the fraction of sequencing reads in the proband carrying the mutated allele, for DNMs inferred to have occurred in either parental germlines (orange) and in the early development of the proband (blue). The expected fraction of ½ for inherited variants is shown as a vertical dashed line. The p-value is obtained from a Kolmogorov-Smirnov test.

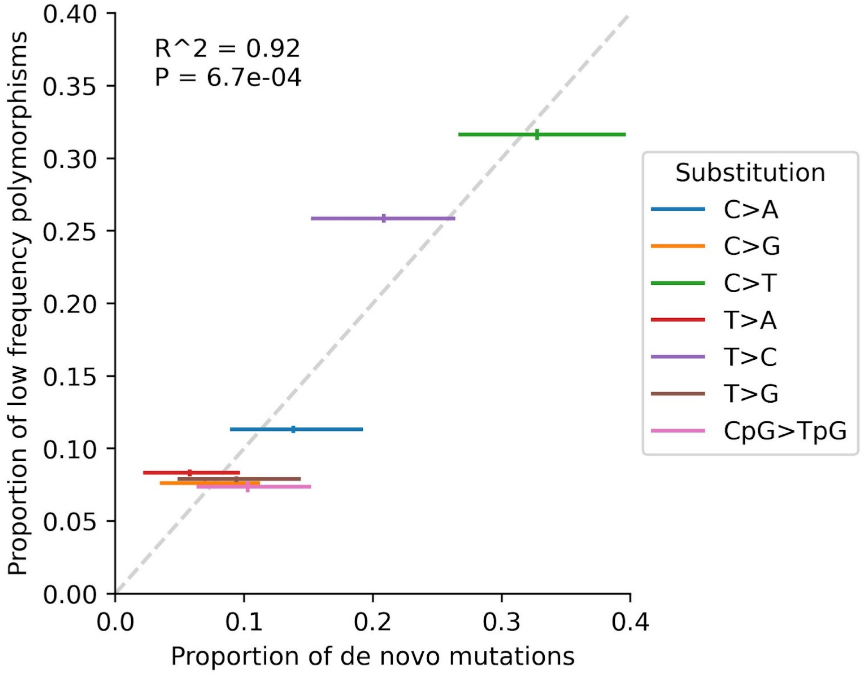


**Figure S3. Proportion of each of seven substitution types.** Proportion of each substitution type in DNMs inferred from pedigree sequencing (x-axis) versus in low-frequency polymorphisms segregating in a dataset of 27 non-closely related individuals (y-axis). 95% CIs (lines) were obtained by bootstrapping 5 Mb autosomal windows 500 times.

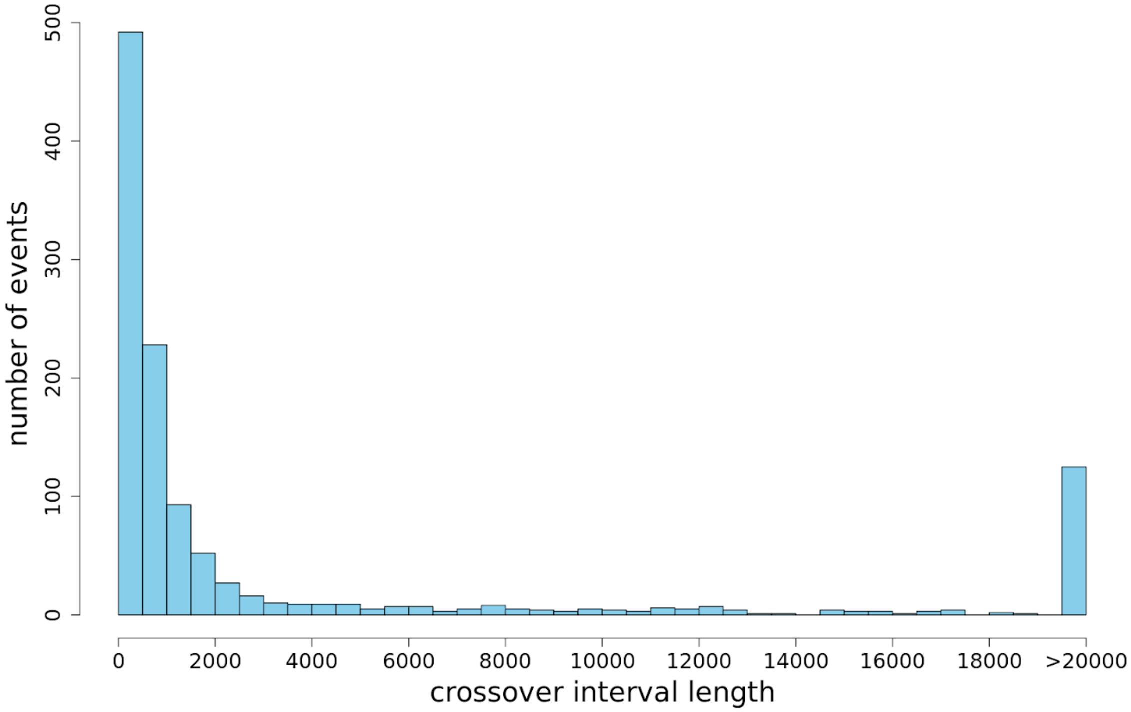


**Figure S4. Distribution of interval lengths within which a crossover is inferred to have occurred.** Intervals larger than 20,000 bps are shown in one bin.

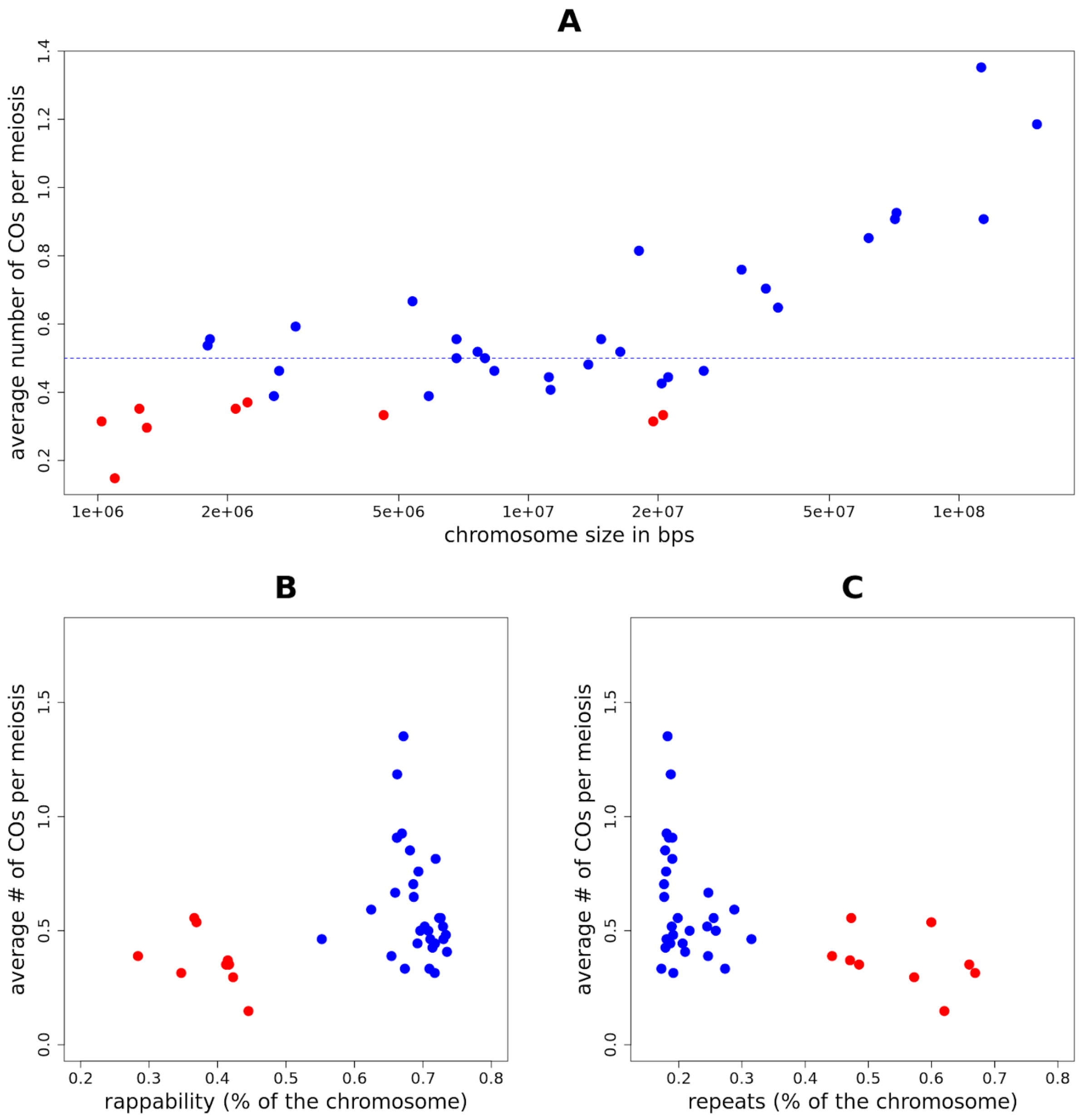


**Figure S5. Crossovers per meiosis as a function of chromosome size in bps, mappability and repeat content for each chromosome.** (A) Average number of crossovers observed per meiosis for the 39 autosomes. The X axis is on a log-scale. The blue dashed line represents the minimum number of crossovers expected per chromosome per meiosis. (B) Average number of crossovers observed per meiosis as a function of the mappability of the chromosome. (C) Average number of crossovers observed per meiosis as a function of the proportion of repeat elements in the chromosome. For each panel, the red dots indicate the chromosomes for which the average number is significantly smaller than 0.5 ($p$ <0.05, using an exact binomial test).

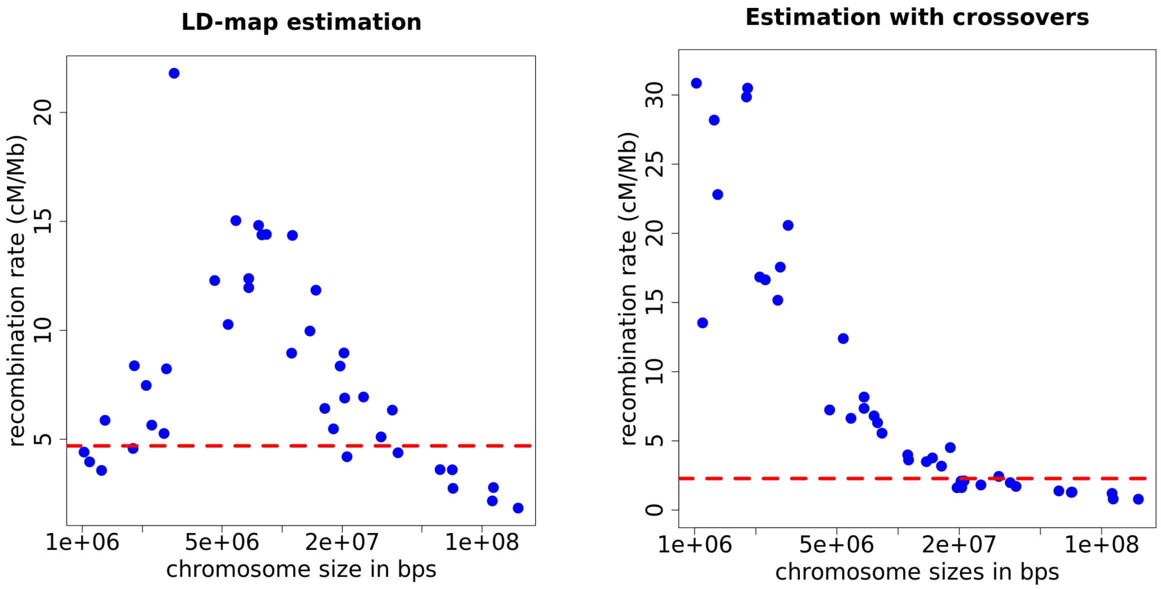


**Figure S6. Mean recombination rate per chromosome.** The left panel shows estimates from the LD-based map; the dashed red line represents the mean recombination rate for the whole genome, including microchromosomes (4.68 cM/MB). The right panel shows estimates from crossovers; the red dashed line is the mean recombination rate for the whole genome is 2.28 cM/Mb. The comparison of these two plots suggests that estimates of recombination rates for chromosomes <40 Mb are unreliable, consistent with simulations (Singhal et al. 2015).

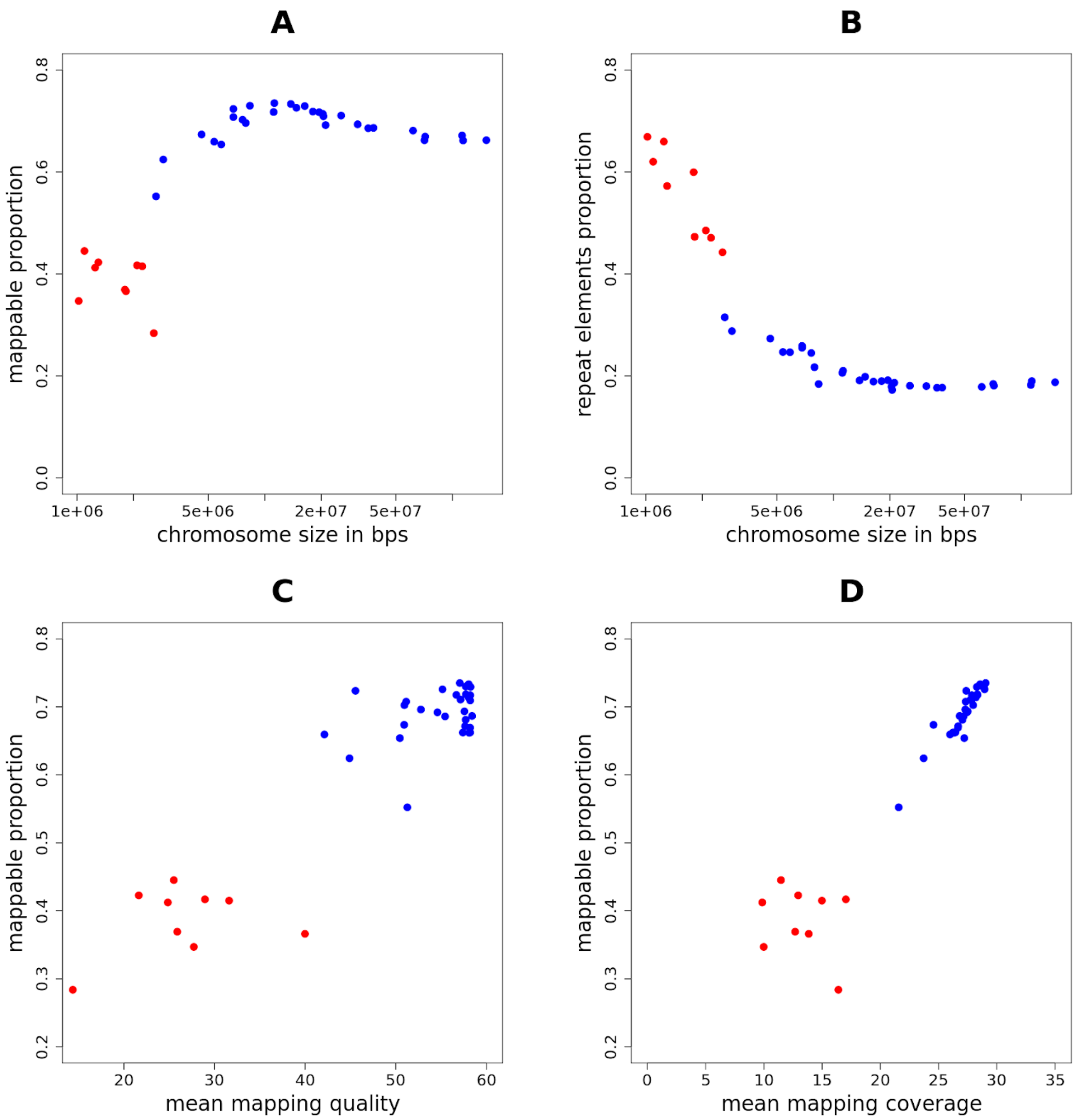


**Figure S7. Proportion of sites that are mappable or repeat elements as a function of chromosome size,mapping quality or coverage.** (A) Fraction of the chromosome that is mappable (see Methods) as a function of chromosome size. (B) Proportion of repeat elements per chromosome as a function of chromosome size. The proportion of repeat elements was estimated by counting the proportion of positions, for each chromosome, that is assigned as a repeat in the softmask version of the genome. (C) Proportion of the chromosome that is mappable as a function of the mean mapping quality for each chromosome. (D) Proportion of the chromosome that is mappable as a function of the mean mapping coverage for each chromosome. For each panel, the red dots are the chromosomes that have been removed from the NCO analysis. The blue dots are the chromosomes that meet our mapping quality and mapping coverage threshold (see Methods).

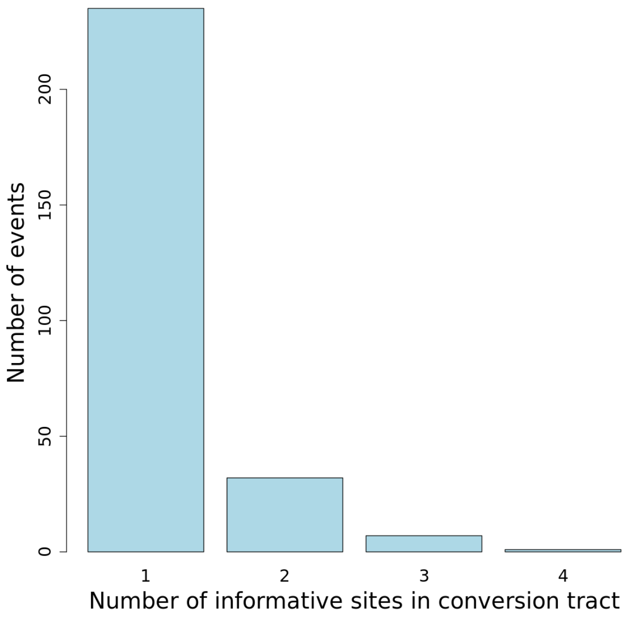


**Figure S8. Distribution of the number of informative sites in the non-crossover conversion tracts.**

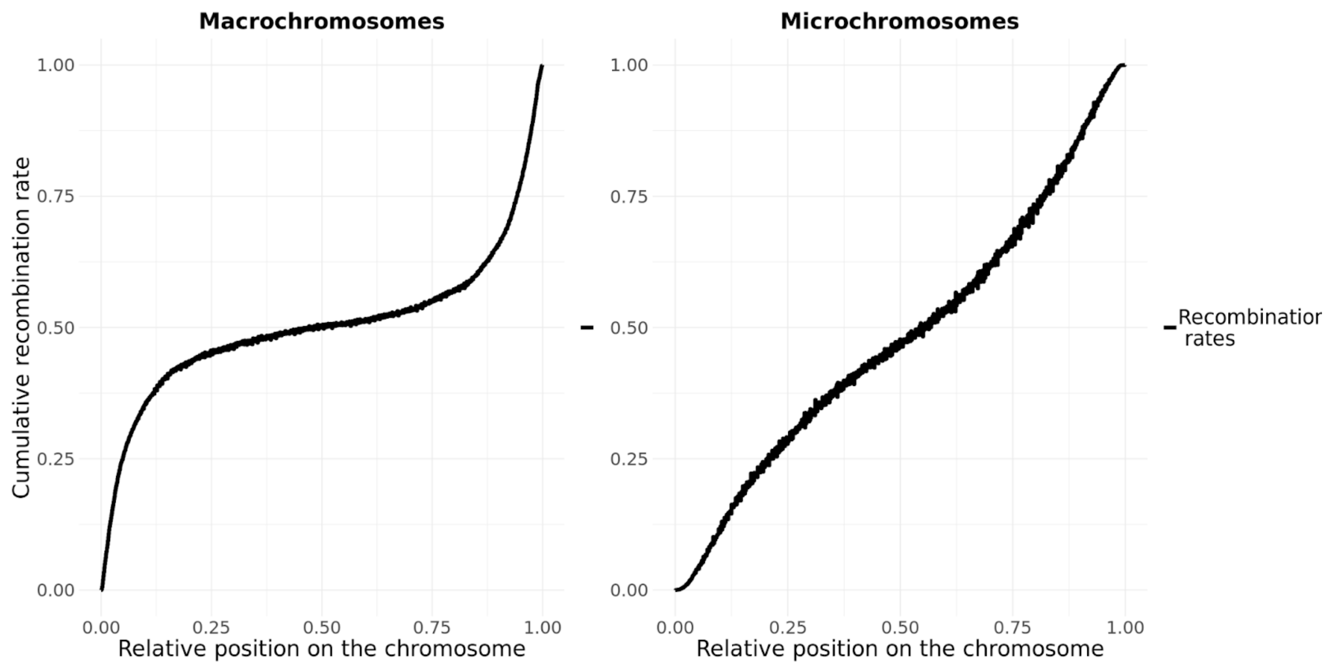


**Figure S9. Cumulative recombination rates for macro- and microchromosomes.** Results are shown for the six macrochromosomes (defined as chromosomes > 40 Mb in size) in the left panel and for the 24 microchromosomes that meet our mapping quality and mapping coverage threshold on the right. The position of events was normalized by the size of the chromosome.

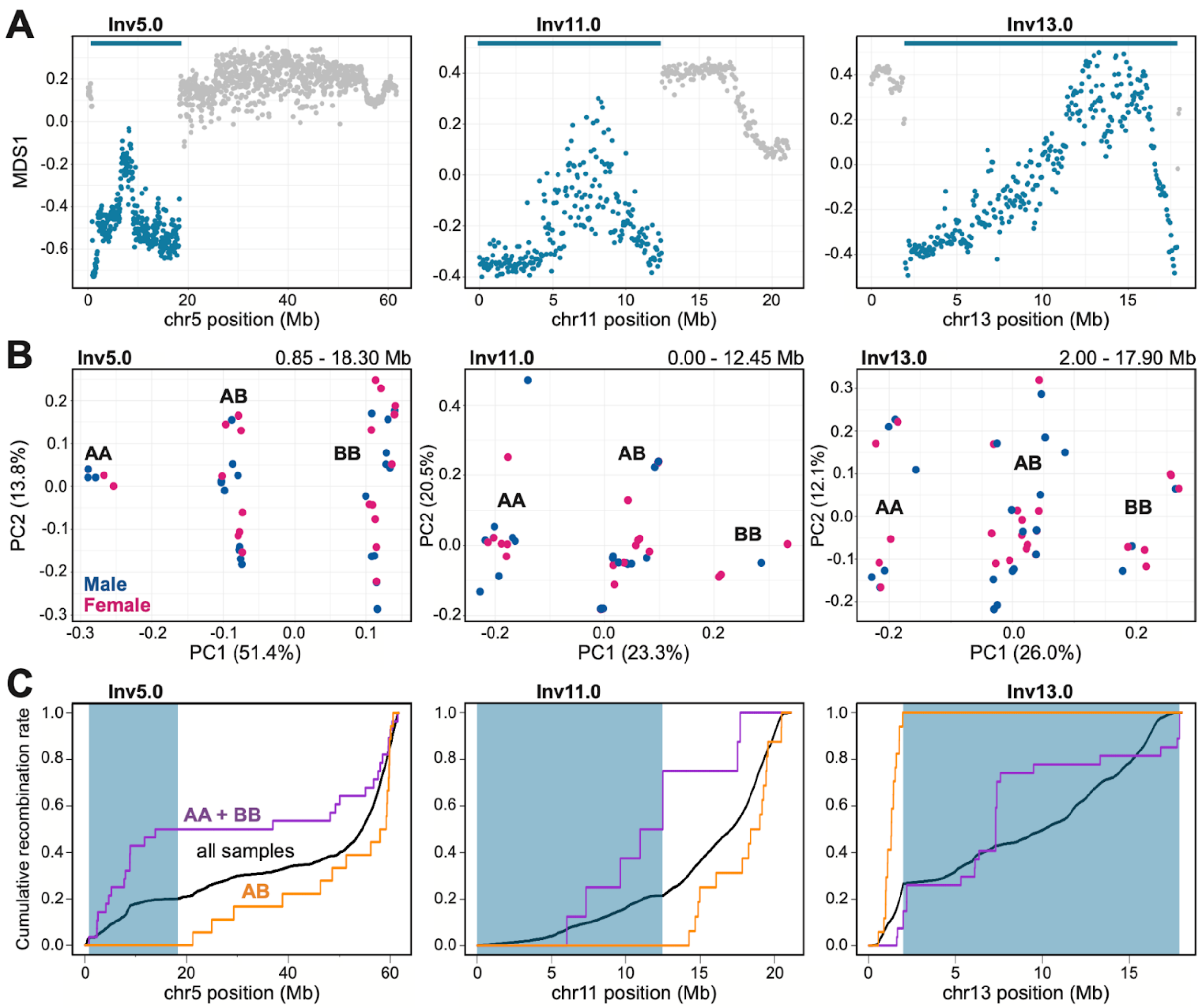


**Figure S10. Chromosomal inversion identification, karyotype assignment, and consequences on crossover distribution and population recombination rate.** (A) Local PCA for three autosomal chromosomes harboring inversion polymorphisms in the zebra finch, with each point representing a 50 kb window. The distance between the relatedness structure of individual windows was evaluated using multidimensional scaling (MDS) and is represented using the MDS1 axis. Windows colored in blue belong to outlier regions associated with the candidate inversions, with the length of each inversion represented as a horizontal blue bar at the top. (B) Consistent with the occurrence of inversion polymorphisms, PCA using variants from the entirety of each outlier region recover three distinct clusters of samples along PC1. These clusters correspond to either group of homokaryotypic (i.e., groups AA and BB) individuals at opposite ends of PC1 and a group of heterokaryotypic individuals (i.e., group AB) immediately in between. Homokaryotype group assignment (i.e., AA vs. BB) was done arbitrarily for each inversion from left to right along PC1 and is not necessarily consistent with labeling in Knief *et al.*, 2016 (Knief *et al.*, 2016). The name of each inversion is given at the top left and the approximate coordinates of each inversion is at the top right of each panel, respectively. Points are color-coded by sex for male (blue) or female (pink). (C) Cumulative distributions of crossovers (purple and orange) and population recombination rate inferred from LD-map (black) for each autosome harboring an inversion polymorphism. Crossovers from homokaryotypic and heterokaryotypic individuals represented in purple and orange, respectively. The location of each inversion is shown in blue.

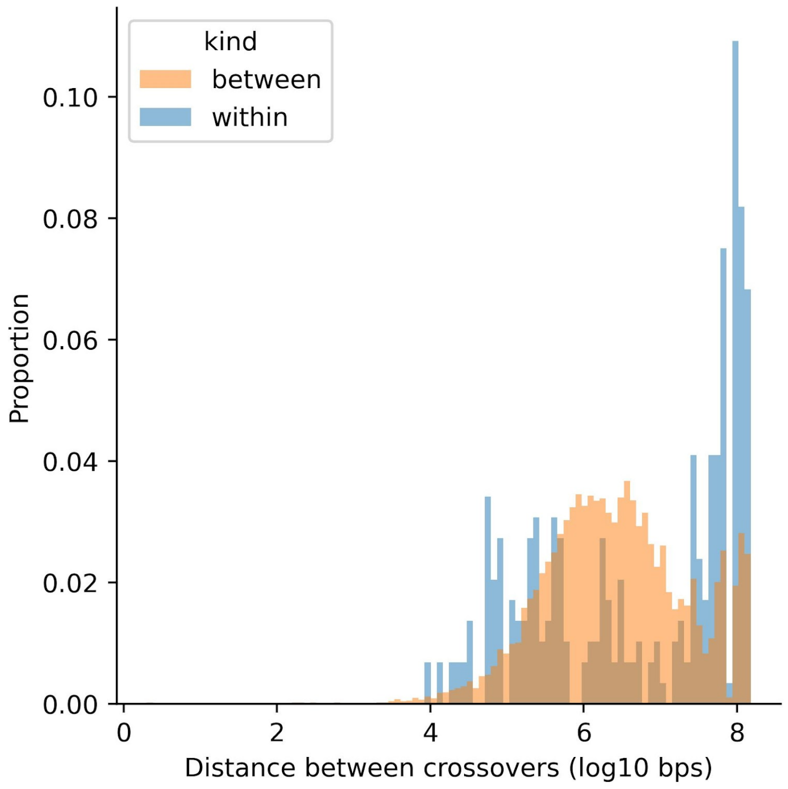


**Figure S11. Distribution of distances between the closest pairs of crossover events, for comparisons of events that occurred within the same meiosis ("within", blue) or a different meiosis ("between", orange).**

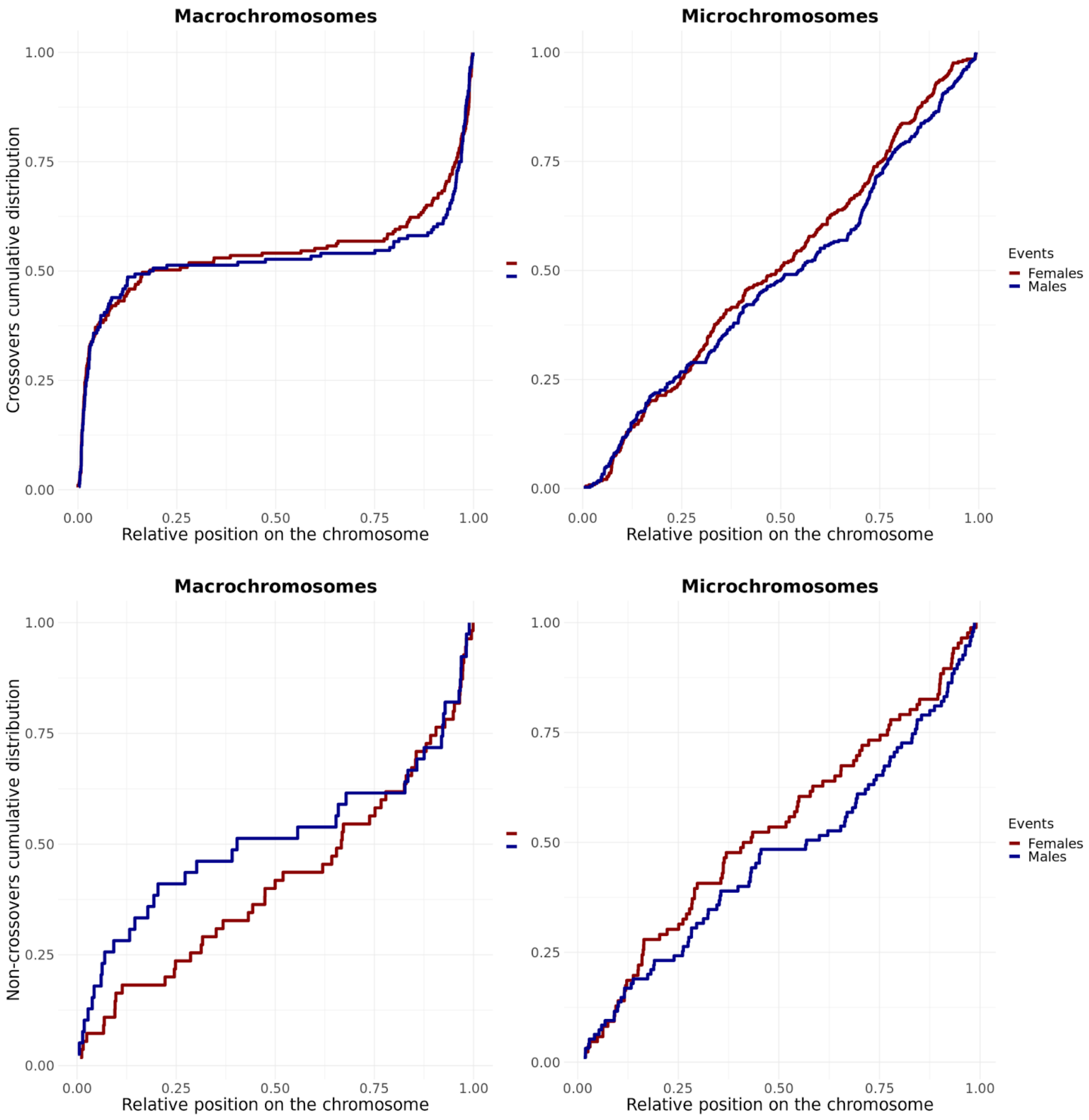


**Figure S12. Cumulative distribution of recombination events for macro- and microchromosomes, in the two sexes.** Top panel: Cumulative distribution of crossover events (females in red and males in blue), for macrochromosomes (left) and microchromosomes (right). The position of the events is normalized by the size of the chromosomes. For crossovers, the *p*-values for Kolmogorov-Smirnov tests comparing the two sexes are 0.34 and 0.22 for the macrochromosomes and the microchromosomes, respectively. Bottom panel: Cumulative distribution of non-crossover events (females in red and males in blue), for macrochromosomes (left) and microchromosomes (right). The position of events is normalized by the size of the chromosomes. For non-crossovers, the *p*-values for Kolmogorov-Smirnov tests comparing the two sexes are 0.15 and 0.32 for the macrochromosomes and the microchromosomes, respectively. For a comparison of the sex-averaged distribution on macro- vs micro-chromosomes, see Table S4.

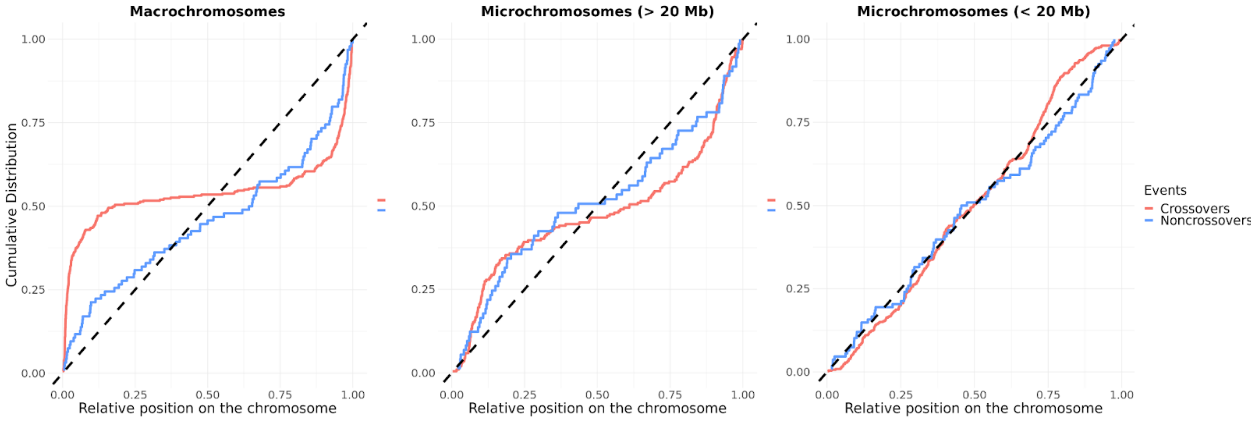


**Figure S13. Cumulative distribution of recombination events with three categories of chromosome sizes.** Same as in Figure 2, but considering chromosomes shorter than 20 Mb separately from those between 20 and 40 Mb and macrochromosomes.

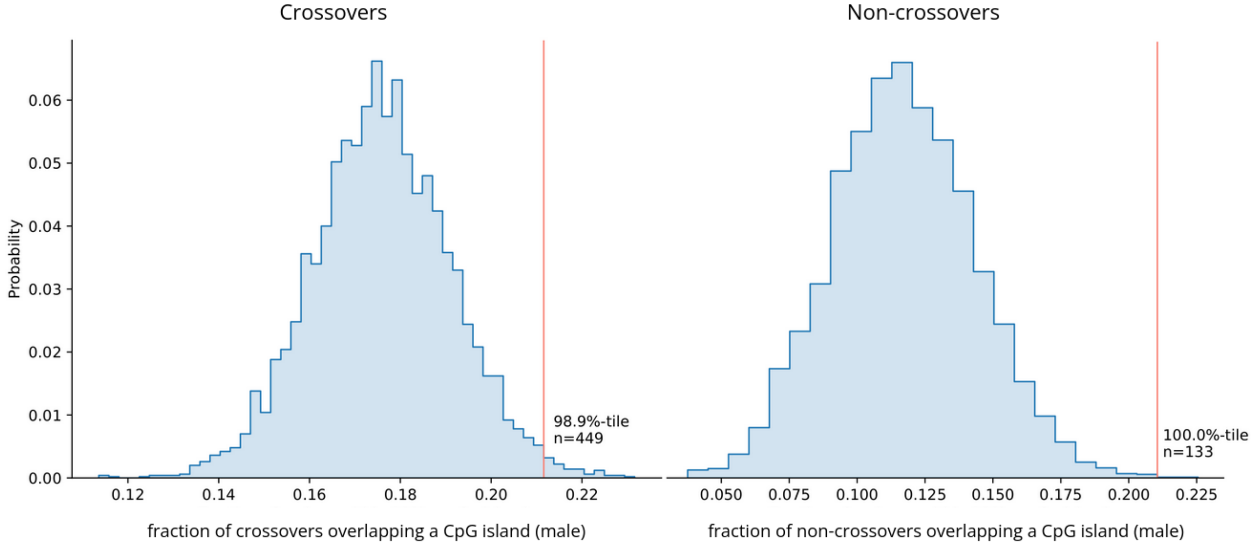


**Figure S14. Overlap of both types of recombination events (in males) with CpG islands.** Fractions of crossovers (top) and non-crossovers (bottom) identified in males meioses within 100 bps of a CpG island. The vertical lines show the observed overlap. The distribution for the overlap expected by chance is shown as a histogram, obtained by randomly shuffling all the events within a 2.5 Mb window on each side of their original location, matching for the GC content and ensuring similar mappability (see Methods for details).

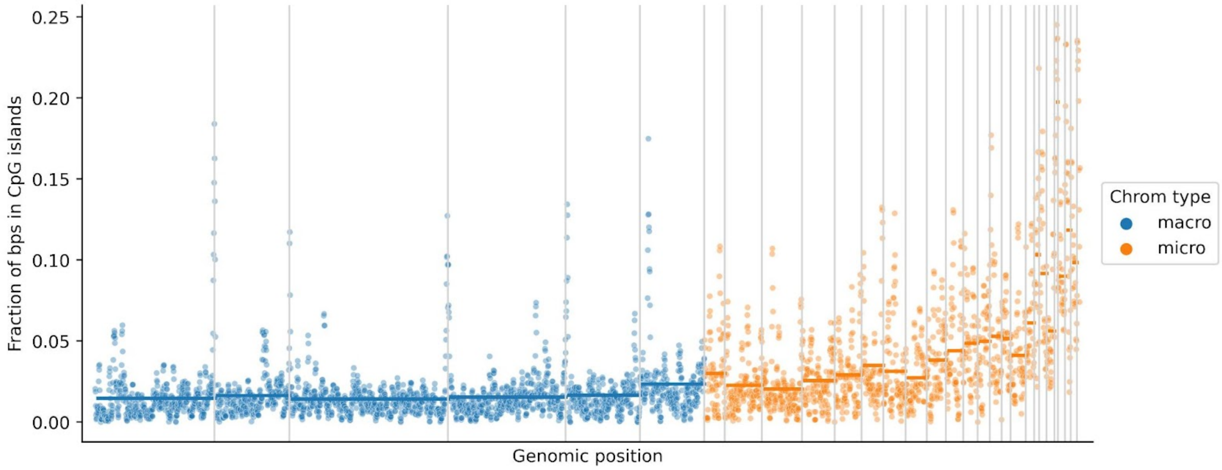


**Figure S15. CpG island density along the zebra finch genome.** The fraction of bps within CpG islands in consecutive genomic windows of 1 Mb, for the 30 autosomes for which we identified recombination events (see Methods). Vertical gray indicates transitions between different chromosomes. Horizontal lines show the mean CpG island density for the corresponding chromosome.

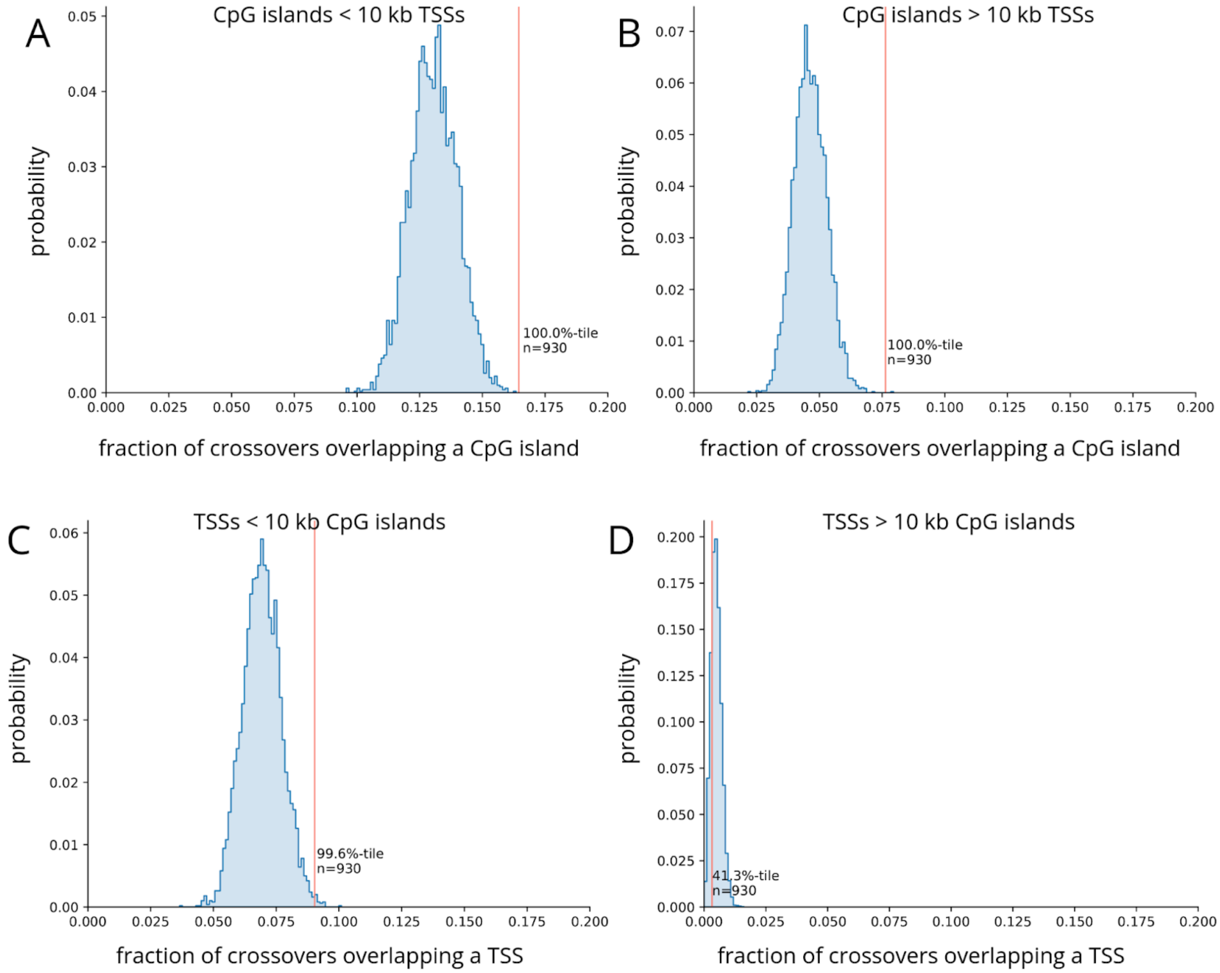


**Figure S16. Overlap of crossovers with CpG islands and transcription start sites (TSSs).** Fraction of crossover events overlapping (**A**) a CpG island at most 10 kb away from a TSS, **(B)** a CpG island farther than 10 kb from a TSS, **(C)** a TSS at most 10 kb away from a CpG island, and **(D)** a TSS farther than 10 kb from a CpG island. To account for the difference in width between CpG islands and TSSs, we considered that a crossover event overlaps a CpG island (A and B) and a TSS (C and D) if it is closer than 100 bps or 500 bps, respectively. The observed overlap is shown by red vertical lines. The overlap expected by chance is shown as a histogram; it was obtained by randomly shuffling all crossovers events 5,000 times within 2.5 Mb of their original location, ensuring the shuffled location had a similar GC content and mappability (see Methods). TSSs were identified from the genome annotation (GCF_003957565.2) by keeping the positions annotated as "start_codon" (n = 20,400).

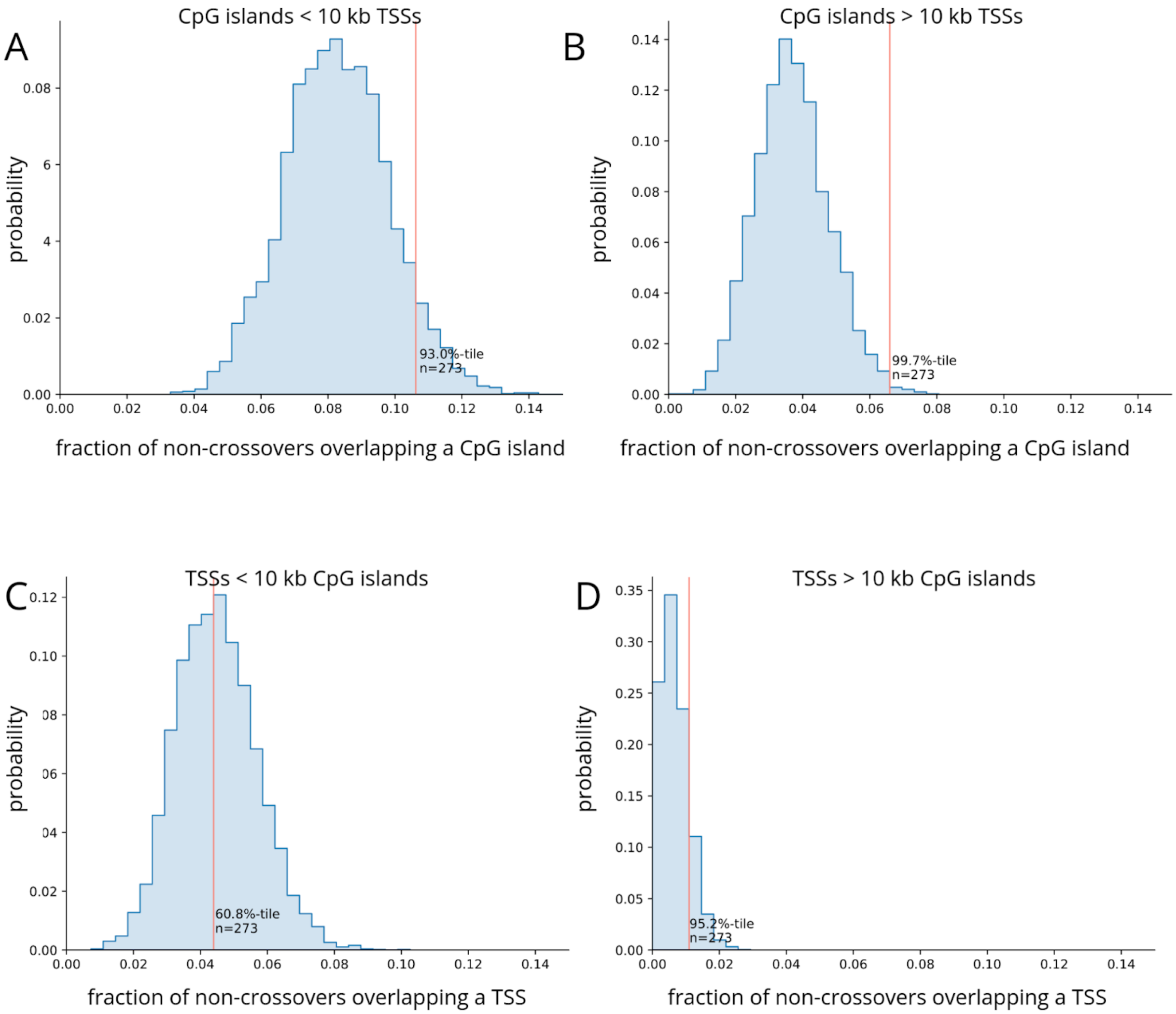


**Figure S17. Overlap of non-crossovers with CpG islands and TSSs.** Fraction of non-crossover events overlapping (**A**) a CpG island at most 10 kb away from a TSS, **(B)** a CpG island farther than 10 kb from a TSS, **(C)** a TSS at most 10 kb away from a CpG island, and **(D)** a TSS farther than 10 kb from a CpG island. To account for the difference in width between CpG islands and TSSs, we considered that a non-crossover event overlaps a CpG island (A and B) and a TSS (C and D) if it is closer than 100 bps or 500 bps, respectively. The observed overlap is shown by red vertical lines. The overlap expected by chance, shown as a histogram, was obtained by randomly shuffling all crossovers events 5,000 times within 2.5 Mb of their original location, ensuring the shuffled location had a similar GC content and mappability (see Methods).

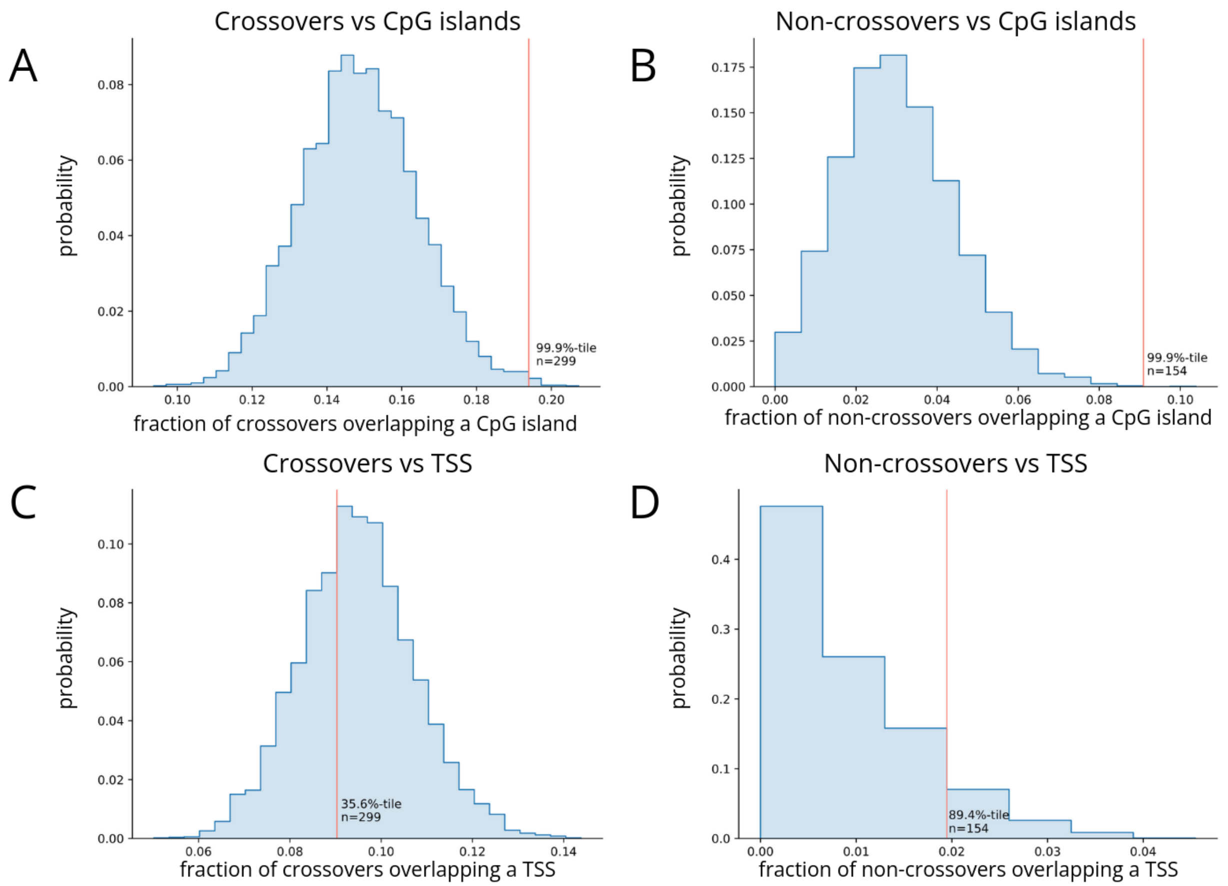


**Figure S18. Overlap of crossovers and non-crossovers with CpG islands or TSSs in collared flycatcher.** **(A)** Fraction of crossovers detected less than 100 bps from a CpG island. **(B)** Fraction of non-crossovers detected less than 100 bps from a CpG island. **(C)** Fraction of crossovers detected less than 100 bps from a TSS. **(D)** Fraction of non-crossovers detected less than 100 bps from a TSS. The vertical lines show the observed overlap. The overlaps expected by chance are shown as histograms, obtained by randomly shuffling 5,000 times all the events within a 2.5 Mb window on each side of their original location, matching for the GC content but not for the mappability (see Methods for details).

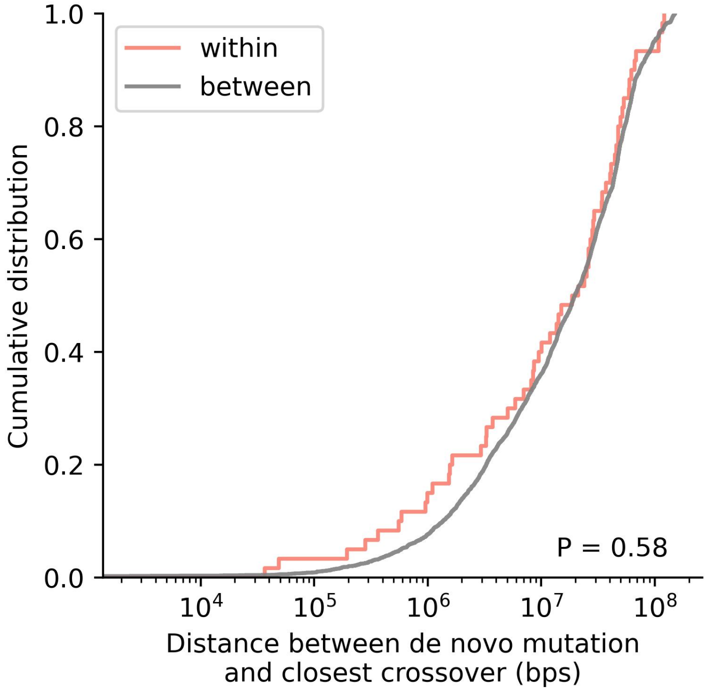


**Figure S19. Cumulative distribution of distances between DNMs and the closest crossover.** In red are the distances between events that occurred in the same germline (i.e., detected in the same proband and assigned to the same parental chromosome) and in gray between events that occurred in different germlines. The p-value was obtained from a Kolmogorov-Smirnov test.

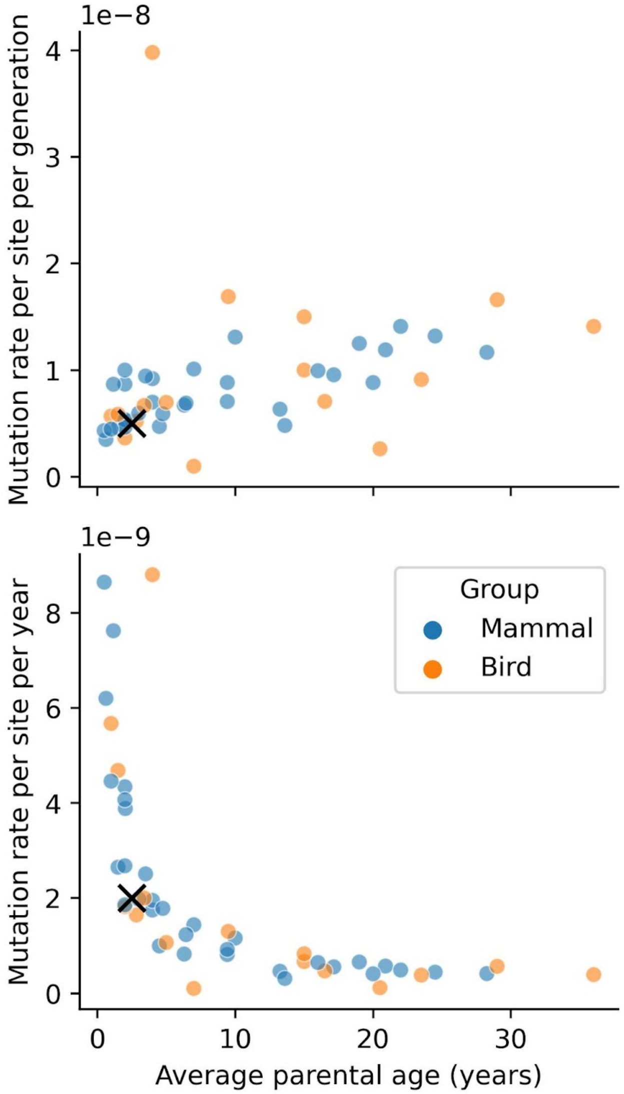


**Figure S20. Mutation rate estimates obtained from pedigree sequencing in vertebrates.** The average mutation rate per generation (top) and per year (bottom) is shown for 48 species of vertebrates. Data from Bergeron et al. 2023 (their Supplementary Table 2); their estimates for mammals and birds are indicated by circles, while our point estimates for zebra finch are represented by crosses.

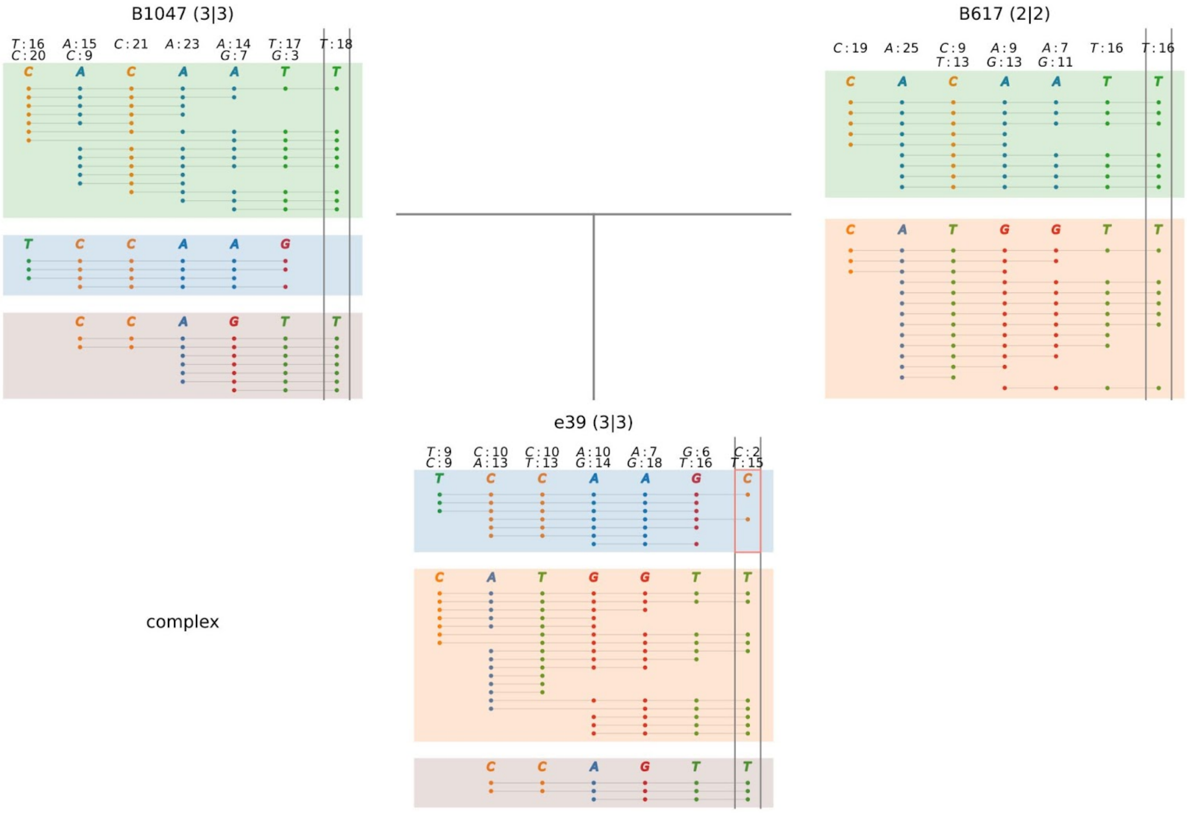


**Figure S21. Example of a region around a putative DNM with evidence of three distinct haplotypes in one of the parents (B1047, top-left panel).** See Figure S1 for more details about how to interpret the figure. Note that the proband has no sequenced descendants, and thus only three individuals (instead of five) are shown.

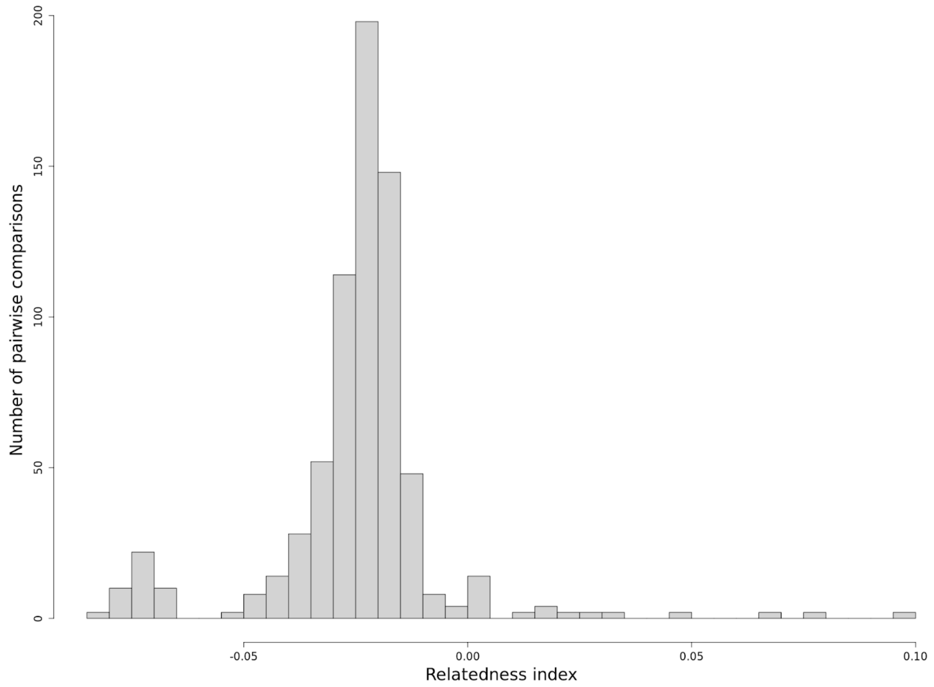


**Figure S22. Histogram of pairwise relatedness for the 27 unrelated individuals.** The relatedness was assessed for each pair of individuals with *relatedness2* tool (VCFtools suite).

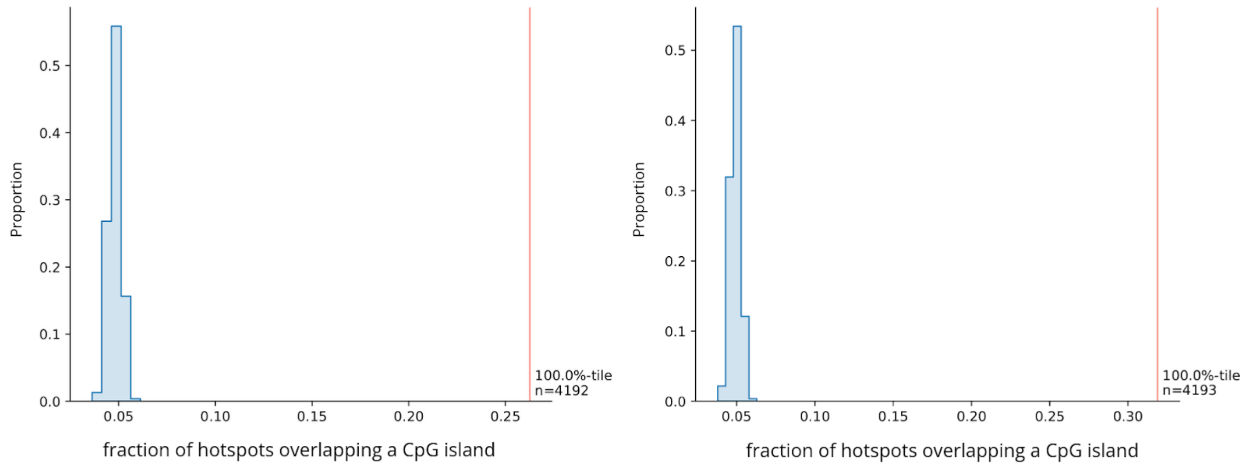


**Figure S23. LD-based hotspots overlap with CpG islands.** Fractions of hotspots within 100 bps of a CpG island, within 10 kb to a TSS (left), or greater than 10 kb from a TSS (right). The vertical red lines show the observed overlap. The overlap expected by chance is shown as a blue histogram, and was obtained by randomly shuffling all the events 5,000 times within a 2.5 Mb window on each side of their original location, matching for the GC content and ensuring a similar mappability (see Methods for details).

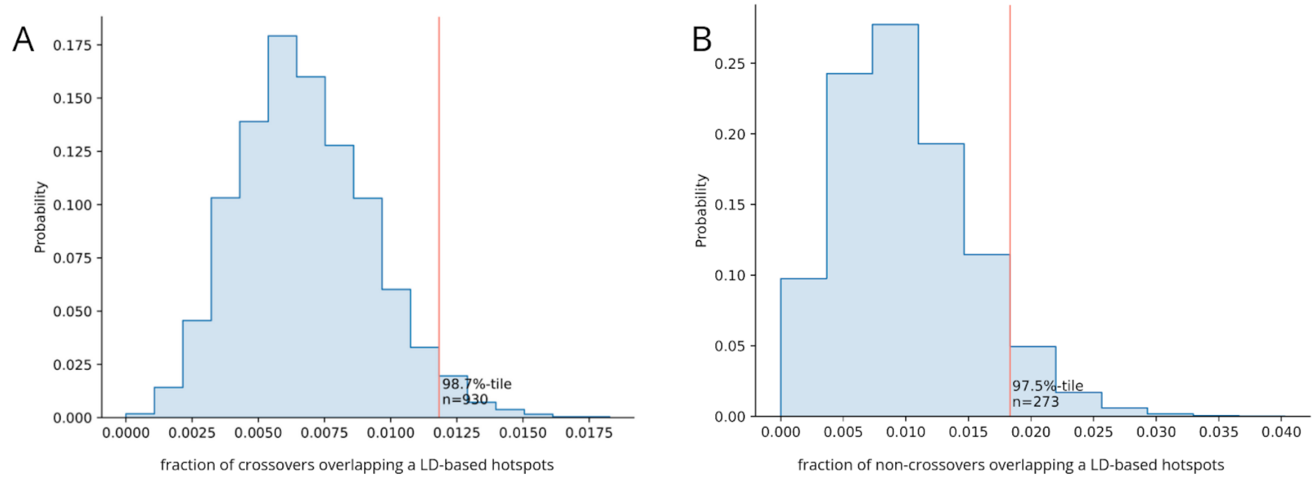


**Figure S24. Recombination events overlap with LD-based hotspots.** Fraction of recombination events within 100 bp from a LD-based hotspot. The overlap expected by chance is shown as a blue histogram, and was obtained by randomly shuffling all the events 5,000 times within a 2.5 Mb window on each side of their original location, matching for the GC content and ensuring a similar mappability (see Methods for details).

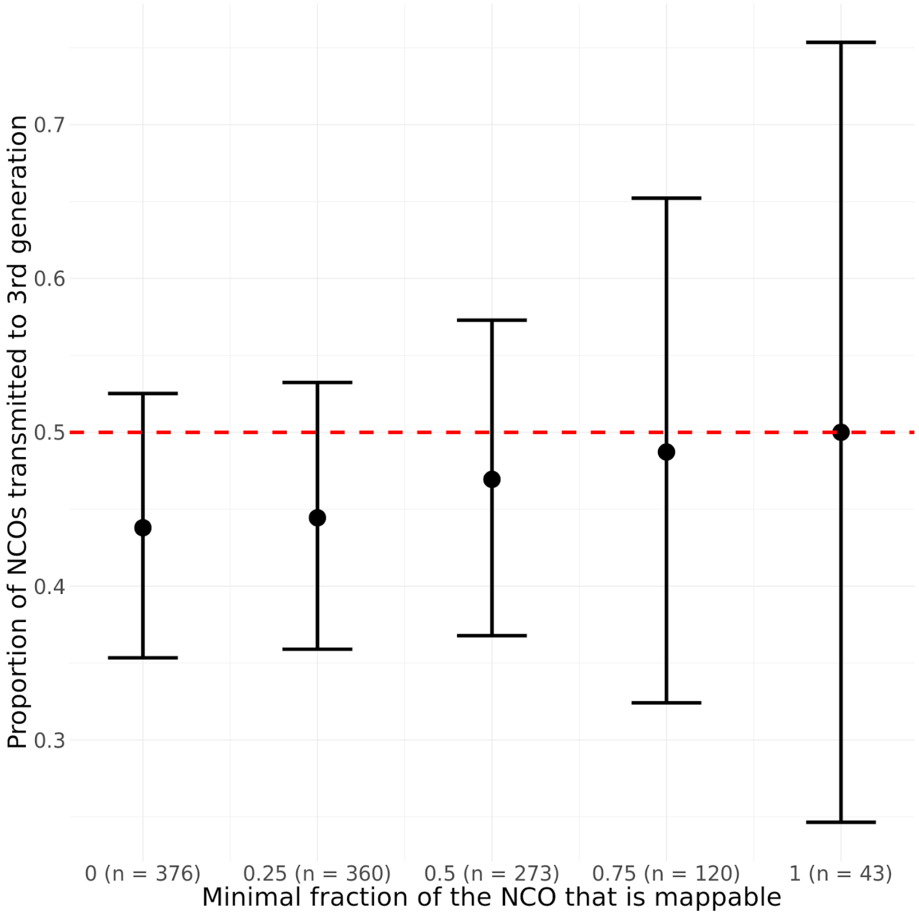


**Figure S25. Transmission rate for different sets of non-crossovers.** The sets were defined depending on the fraction of the length of the non-crossover interval that is mappable in the reference genome (see Method section "Detection of non-crossover events"). The number of non-crossovers in each set is in parentheses on the x axis label. The 95% confidence interval is represented by the black bars. The dashed red line denotes the expectation of 0.5 (assuming perfect power to detect a heterozygous site).

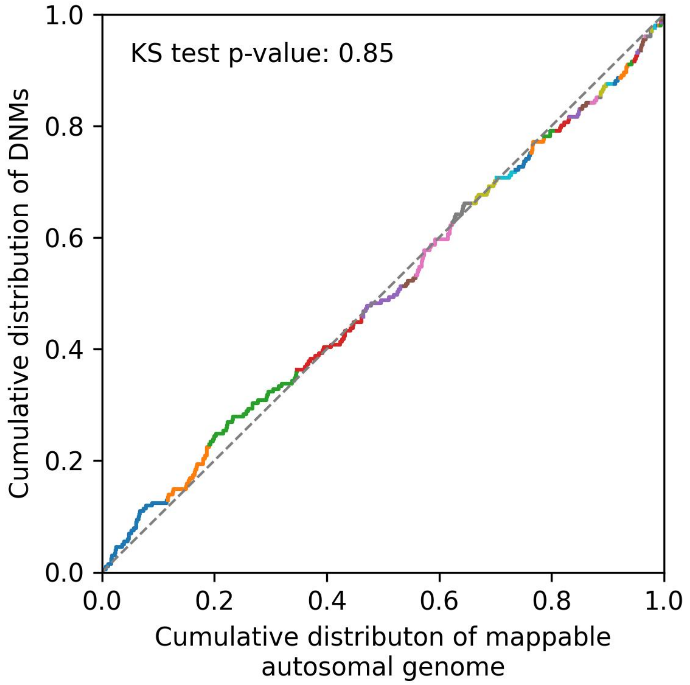


**Figure S26. Distribution of DNMs along the mappable autosomal regions. The p-value of a Kolmogorov-Smirnov test against a uniform distribution is annotated.**

**Table S4. Numbers of recombination events identified in maternal and paternal meioses.** The p-value was obtained from an exact binomial test of no sex difference). For the crossovers, the number of maternal meioses is higher than paternal meioses, and we adjusted the null model accordingly.

| | Crossovers | Non-crossovers |
|---|---|---|
| Maternal | 559 | 141 |
| Paternal | 520 | 134 |
| Total | 1,079 | 275 |
| *p*-value | 1 | 0.72 |

**Table S5. Non-crossover transmission rates for three different variant callers.** Transmission is tested only for configurations in which the genotypes of the offspring, the partner and the grandkid allow us to track the conversion event (see Methods). Shown as CIs are the 95% confidence intervals from an exact binomial test.

| Variant caller | # NCOs | Testable | Transmitted | Rate | CI |
|---|---|---|---|---|---|
| *GATK* | 121 | 56 | 21 | 0.38 | 0.25 - 0.51 |
| *bcftools* | 288 | 96 | 40 | 0.42 | 0.32 - 0.52 |
| *freebayes* | 273 | 99 | 47 | 0.47 | 0.37 - 0.57 |

**Table S6. *P*-values for the Kolmogorov-Smirnov tests presented in Figure 2.** For each set of chromosomes, we tested if the distribution of crossovers is the same as that of non-crossovers, and whether the distribution of each type of recombination events are uniform. We also compared the distribution of crossovers (non-crossovers) on macro- vs microchromosomes

| **Chromosomes tested** | **Recombination events tested** | ***p*-values** |
|---|---|---|
| **On macrochromosomes** | | |
| | Crossovers *vs* non-crossovers | $7.0 \times 10^{-5}$ |
| | Crossovers uniform? | $< 2.2\ 10^{-16}$ |
| | Non-crossovers uniform? | $9.1 \times 10^{-5}$ |
| **On microchromosomes** | | |
| | Crossovers *vs* non-crossovers | 0.38 |
| | Crossovers uniform? | 0.006 |
| | Non-crossovers uniform? | 0.23 |
| **Using all chromosomes** | | |
| | Crossovers on macro- *vs* microchromosomes | $< 2.2\ 10^{-16}$ |
| | Non-crossovers on macro *vs* microchromosomes | 0.16 |

**Table S7. Average number of pairs of phase changes per family per meiosis.**

| Pedigree | pairs of phase changes |
|---|---|
| Family 1 | 17.8 |
| Family 2 | 12.0 |
| Family 3 | 18.6 |
| Family 4 | 41.0 |

**Command lines**

Representative pseudo-code lines to reproduce the analyses. The more detailed scripts and READMEs can be found at https://doi.org/10.5281/zenodo.13696268.

| Whole genome sequencing alignment |
| --- |
| gatk HaplotypeCaller -R {ref} -I {bam} -O {vcf} --heterozygosity 0.01 \ <br> -ERC BP_RESOLUTION -L {chrom} --pcr-indel-model NONE <br> gatk GenomicsDBImport --sample-name-map {input} -R {ref} -L {chrom} \ <br> --genomicsdb-workspace-path {output} <br> gatk GenotypeGVCFs -R {ref} -V gendb://{input} -O {output} --heterozygosity 0.01 \ <br> -L {chrom} |

| GATK variant calling |
| --- |
| gatk HaplotypeCaller -R {ref} -I {bam} -O {vcf} --heterozygosity 0.01 \ <br> -ERC BP_RESOLUTION -L {chrom} --pcr-indel-model NONE <br> gatk GenomicsDBImport --sample-name-map {input} -R {ref} -L {chrom} \ <br> --genomicsdb-workspace-path {output} <br> gatk GenotypeGVCFs -R {ref} -V gendb://{input} -O {output} --heterozygosity 0.01 \ <br> -L {chrom} |

| Generation of mappability mask |
| --- |
| splitfa {Fasta} 150 | split -l 20000000 --filter='gzip > \$FILE.gz'- kmers/{Chunk} <br><br> bwa aln -R 1000000 -O 3 -E 3 {fasta} kmers/{Chunk} > bwa/{Chunk}.sai <br><br> bwa samse {Fasta} bwa/{Chunk}.sai kmers/{Chunk} | gzip > bwa/{Chunk}.sam.gz <br><br> gzip -dc bwa/*.sam.gz | perl gen_raw_mask.pl > rawMask_150.fa <br><br> gen_mask -l 150 -r 0.5 rawMask_150.fa > mask_150.fa <br><br> python makeMappabilityMask.py mask_150.fa > {Mappability_mask.bed} |

#### Detection of single-point *de novo* mutations

```
python scripts/call_dnms.py -i {vcf} -t {trio} -c {chrom}
```

#### Assigning parent of origin of *de novo* mutations

```
python scripts/classify_dnms.py {bams} -v {vcf} -m {start}
```

#### Inference of mutation signature activity

```
Analyze.cosmic_fit({input_matrix}, {output_dir},  input_type="matrix")
```

#### Population structure and relatedness

```
vcftools --gzvcf {input.vcf} --chrom-map {input.chrommap} --plink --out {params.outname1} --chr {wildcards.chromosome} --temp tmp/

plink --allow-extra-chr --geno 0.10 --chr-set 1 no-xy no-mt --indep-pairwise 50 5 0.2 --maf 0.05 --out {params.outname2} --file {params.outname1}

plink --chr-set 1 no-xy no-mt --allow-extra-chr --pca --file {params.outname1} --extract {params.outname2}.prune.in --out {params.outname3}

zcat {input.vcf} | vcfsnps | vcfbiallelic | vcftools --vcf - --weir-fst-pop {input.pop1} --weir-fst-pop {input.pop2subset} --fst-window-size {params.wind_size} --fst-window-step {params.wind_step} --out {params.outfst_sub}

zcat {input.vcf} | vcfsnps | vcfbiallelic | vcftools --vcf - --relatedness2 --out {params.outrelatedness} --exclude-bed {params.toexlude}
```

#### Diversity and effective population size estimates

```
bedtools coverage -sorted -a {input.mappable} -b <(bcftools filter -g 5 {input.vcf} | vcfsnps) > {output}
```

### Ancestral alleles and mutation transition matrix

```
#### test if allele is significantly major

        vcftools --gzvcf {input} --freq --out out_ancestral/onesided_test_unrelated/{wildcards.chromosome}
        sed '1d' out_ancestral/onesided_test_unrelated/{wildcards.chromosome}.frq -i
        python scripts/binom_test_major_allele_exceed.py
    out_ancestral/onesided_test_unrelated/{wildcards.chromosome}.frq {output.all}
        awk 'length($8)==1 && length($9)==1' {output.all} > {output.all}_tmp
        mv {output.all}_tmp {output.all} -f
        awk '{{if ($7<0.05) print}}' {output.all} > {output.filtered}

#### compute mutation transition matrix

    cat  {output.filtered} > {params.mut}
    python scripts_compute_mut_mat.py {params.mut} {input.fasta} {output}
```

### Variant phasing

```
    gatk VariantFiltration -V {input} -O {output.tmp} "
        "--filter-expression 'QD < 6.0 || MQ < 40.0 || FS > 10.0 || SOR > 4.0 || "
        "MQRankSum < -12.5 || ReadPosRankSum < -8.0' --filter-name 'hard_filter' --verbosity ERROR;"
        "vcftools --gzvcf {output.tmp} --remove-filtered-all --max-alleles 2 "
        "--recode --recode-INFO-all --stdout | bgzip > {output.vcf};"
        "tabix -f -p vcf {output.vcf}

    bcftools filter -g 5 {input} | vcfsnps | bgzip > {output.vcf}
        tabix -f -p vcf {output.vcf}

    extractPIRs --bam {params.bam_list} --vcf {input} --out {output.outfile}

    plink2 --allow-extra-chr --update-parents {input.ped_info} --vcf {input.vcf} --make-bed --out {params.out_name}
```

```
shapeit -assemble --input-bed {params.bed} --aligned --input-pir {input.PIRs} -O {params.out} --effective-size {params.N} --rho {params.rho} --duohmm -W {params.windows} --output-log {params.log_dir}/{wildcards.chromosome} --force --thread {params.num_threads}

shapeit -convert --aligned --input-haps {params.file_in} --output-vcf {output} --output-log {params.log_dir}/{wildcards.chromosome}

vcftools --ldhelmet --vcf {input} --chr {wildcards.chromosome} --out vcftools_output/{wildcards.chromosome} --keep {params.tokeep} --temp {params.tmpdir}
```

### LDhelmet analysis

```
ldhelmet find_confs --num_threads {params.cpus} -w {params.windows} -o {output.confs} {input}

ldhelmet table_gen --num_threads {params.cpus} -c {input.confs} -t {params.theta} -r 0.0 .0000001 .000001 .000001 .00001 .00001 .0001 .0001 .001 .001 .01 .01 .1 .1 1.0 0.5 50.0 -o {output}

ldhelmet pade --num_threads {params.cpus} -c {input.confs} -t {params.theta} -x {params.coeff} -o {output}

ldhelmet rjmcmc --num_threads {params.cpus} -w {params.windows} -l {input.likelihoods} -p {input.pade} -b {wildcards.bkpty} --snps_file {input.haplotypes} -m {input.mutmat} --burn_in {params.burn_in} -a {input.ancestral} --pos_file {input.positions} -n {params.iterations} -o {output}

ldhelmet post_to_text -m -p {params.quantile_1} -p {params.quantile_2} -p {params.quantile_3} -o {output} {input}
```

### Identification of hotspots

```
bedtools intersect -a  {input.genome} -b {input.rec} -wo | compute_rrate_windows.sh > {output}

bedtools map -a {input.windows} -b {input.recrate} -c 4 -o mean > {params.meanrate}
python scripts/relative_recomb_rate.py {params.meanrate} > {output}
```

### Identification of CpG islands

```
cpgplot -sequence ${REF} -outfile ${CPGPLOT_OUT} -window 50 -minlen 250 \\
  -minoe 0.6 -minpc 50 -graph png -outfeat ${CPGI.GFF} -plot No
python scripts/HMM.py -s {chrom}
```

#### Detection of crossovers

```
python scripts/call_crossovers.py  -i {vcf} -f {fam_ped} -m {min_info_sites} -p {out_pdf} -v {prefix} -o {out_bed}
```

#### *Freebayes* variant calling

```
bamaddrg {bams} -R {region} -c \
 | freebayes -f {ref} --stdin -g 10000 --min-mapping-quality 30 \
   --min-base-quality 30 --use-best-n-alleles 4 \
 | vcfallelicprimitives -kg \
 | bgzip > {vcf}
```

#### *Bcftools* variant calling

```
bcftools mpileup -b {bams} -f {ref} --max-depth 150 --min-BQ 30 \
   --min-MQ 30 -r {region} -a AD,ADF,ADR,SP,INFO/AD,INFO/ADF,INFO/ADR -Ou \
 | bcftools call -Ou -mv \
 | bcftools filter -s LowQual -e '%QUAL<20' -Oz -o {vcf}
```

#### Detection of non-crossovers events (including extra filtering steps)

```
zcat {input.vcf} | grep -v "\.:\.:\.:\."  | python {params.script1} -s {input.depth} -u {params.upper} -l {params.lower} | bgzip > {params.tmpfile}

bcftools query {input.vcf} -f '%CHROM\t%POS\t%REF\t%ALT\t[%SAMPLE=%GT\t%RO\t%DP\t%AD\t]\n' -S {input.GdOff_ind} | grep "=\." -v | python3.9 {params.script2} | awk '{{print $1"\\t"$2"\\t"$2}}' | uniq | bgzip > {params.postoremove}

bcftools view -T ^{params.postoremove} -o {output} {params.tmpfile} -O z
```

```
bcftools +mendelian {input.vcf}  -T {input.trio} -d | bgzip > {output.outvcf}

bcftools query -S {input.gd_off} {input.vcf} -f '%CHROM\t%POS\t%REF\t%ALT\t[%SAMPLE=%GT\t%RO\t%DP\t%AD\t]\n' | python {params.script} | grep "\./\." -v | cut -f1-2 > {params.tmp_pos}

tabix -h {input.vcf}  -R <(grep {wildcards.chromosome} {input.mappable} | awk '{{if ($2==0) print $1"\\t""1""\\t"$3; else print $1"\\t"$2"\\t"$3}}') | bgzip > {output.outvcf}

python scripts_noncrossovers_detection.py {input.ped_info} {input.vcf} {output.maternal} {output.paternal} {params.maternal_dir} {params.paternal_dir}
```

#### Phasing of non-crossovers

```
whatshap  phase --reference {ref} --ped {ped} --use-ped-samples {vcf} {bams}  -o {output}
```

#### Conversion tract length estimation

```
Rscript script_lileklyhood.R {params.dir_in} {output}
```

#### Estimating the number of non-crossovers in a meiosis

```
# R code to sample 10,000 windows from the exponential distribution of rate 1/15
n <- 10000

rate_15bp <- (1/15)
samples_15bp <- rexp(n, rate_15bp)
# head(samples)

rounded_samples_15bp <- round(samples_15bp)
write.table(rounded_samples_15bp, file = "sampling_from_exp_distrib_15bp.txt", row.names = FALSE, col.names = FALSE)
```

```
## bash loop for 100 iterations

for i in {{1..100}}
    do
    cat {input.sampling} | awk '{{print "{wildcards.chromosome}""\\t""1""\\t"$1+1}}' | bedtools shuffle -i - -g
{input.chr_size} -chrom | \
    bedtools sort | uniq | bedtools intersect -a - -b {input.info_sites} -wao | awk '{{if ($1==$4) print}}' | cut -f1-3 | uniq
-c | awk '{{print $2"\\t"$3"\\t"$4"\\t"$1}}' > {params.suffix}_${{i}}.bed
     done
```
